# Protein language model-guided engineering of an anti-CRISPR protein for precise genome editing in human cells

**DOI:** 10.1101/2023.12.13.571376

**Authors:** Julia Marsiglia, Kia Vaalavirta, Estefany Knight, Muneaki Nakamura, Le Cong, Nicholas W. Hughes

**Affiliations:** Acrobat Genomics Inc., San Carlos, CA 94070; Department of Pathology, Stanford University School of Medicine, Stanford, CA, 94035, USA; Department of Genetics, Stanford University School of Medicine, Stanford, CA, 94035, USA

## Abstract

Promiscuous editing by CRISPR/Cas systems within the human genome is a major challenge that must be addressed prior to applying these systems therapeutically. In bacteria, CRISPR/Cas systems have evolved in a co-evolutionary arms race with infectious phage viruses that contain inhibitory anti-CRISPR proteins within their genomes. Here, we harness the outcome of this co-evolutionary arms race to engineer an AcrIIA4 anti-CRISPR protein to increase the precision of CRISPR/Cas-based genome targeting. We developed an approach that specifically leveraged (1) protein language models, (2) deep mutational scanning, and (3) highly parallel DNA repair measurements within human cells. In a single experiment, ∼10,000 AcrIIA4 variants were tested to identify lead AcrIIA4 variants that eliminated detectable off-target editing events while retaining on-target activity. The candidates were further tested in a focused round of screening that included a high-fidelity version of Cas9 as a benchmark. Finally, arrayed experiments using Cas9 delivered as ribonucleoprotein were conducted that demonstrated an increase in gene editing precision across two independent genomic loci and a reduction in the frequency of translocation events between an on-target and off-target site. Thus, language-model-guided high-throughput screening is an effective way to efficiently engineer AcrIIA4 to increase gene editing precision, which could be used to improve the fidelity of gene editing-based therapeutics and to reduce genotoxicity.

## INTRODUCTION

CRISPR/Cas technologies^1–3^ have emerged as powerful tools to interrogate gene function at genome-scale in the laboratory^4,5^ and as potentially curative therapeutics^6,7^. CRISPR/Cas9 found in *Streptococcus pyogenes* (SpCas9) relies on single-guide RNA (sgRNA) molecules to direct it to specific genomic loci inside the nucleus. While this mechanism enables programmable editing due to the modularity of the sgRNA sequence, partial complementarity between the sgRNA and multiple genomic loci in the same cell can lead to unintended nuclease activity and genotoxicity^8^. This is of particular concern in a clinical setting where, for example, an unintended edit within a tumor suppressor gene or a chromosomal translocation could lead to oncogenesis^9,10^. Additionally, human genetic variation can create off-target sites that are not present in the reference genome^11^. Therefore, there have been extensive efforts to mitigate off-target effects through (1) computational predictions of off-target regions based on the sgRNA sequence^12,13^ and (2) engineering of the sgRNA and Cas nuclease components^14–17^.

These approaches have been effective but require costly engineering campaigns for each independent CRISPR/Cas ortholog and serial optimization of sgRNAs. As a result, there is demand for additional approaches, using orthogonal components, to increase gene editing precision. The native activity of anti-CRISPR (Acr) proteins to inhibit CRISPR/Cas nuclease activity has been leveraged to increase gene editing precision through re-dosing of AcrIIA4 after Cas9 RNP delivery^18^. Another study analyzed whether fusion of three AcrIIA4 protein variants that have attenuated inhibitory activity with SpCas9 could increase gene editing precision^19^. We sought to build upon this conceptual framework to explore whether further high-throughput engineering of the interaction between AcrIIA4 and SpCas9 could be harnessed to fine-tune gene editing precision.

Computationally guided protein design has emerged as a powerful approach to engineer proteins with desirable properties^20,21^. We wondered whether this approach, coupled with a classic structure-guided engineering approach, could identify AcrIIA4 variants that enable precise genome editing outcomes. We hypothesized that identification of engineering hotspots, which are predicted to have a strong functional impact on AcrIIA4 function, followed by deep mutational scanning of these regions would be a cost-effective strategy to navigate the vast amino acid search space to identify optimal variants. Here, we developed an integrated computational and experimental approach to screen ∼10,000 AcrIIA4 variants in human cells expressing SpCas9 and validated the activity of lead variants, which eliminated detectable off-target activity in the pooled screen setting and within subsequent benchmarking experiments. Together, these results established a general framework that can be further applied to other engineering challenges.

## RESULTS

### Evolving the AcrIIA4 interaction with SpCas9 to increase gene editing precision using PLM-guided library design

We aimed to tune the gene editing activity of CRISPR/Cas9 by engineering the interaction between it and its cognate anti-CRISPR protein, AcrIIA4. By attenuating the activity of AcrIIA4, we hypothesized that we could inhibit off-target activity by CRISPR/Cas9 while retaining on-target activity. We also surmised the interaction between AcrIIA4 and SpCas9 could be evolved using a pooled protein screening approach to reduce off-target activity without the requirement for precisely timed re-dosing of the AcrIIA4 component. More specifically, through protein-language-guided selection of AcrIIA4 variants and pooled protein screening, we aimed to reduce the affinity of AcrIIA4 to SpCas9 and allow for on-target activity while minimizing the potential for off-target activity (**Figure 1A)**.

**Figure 1.**
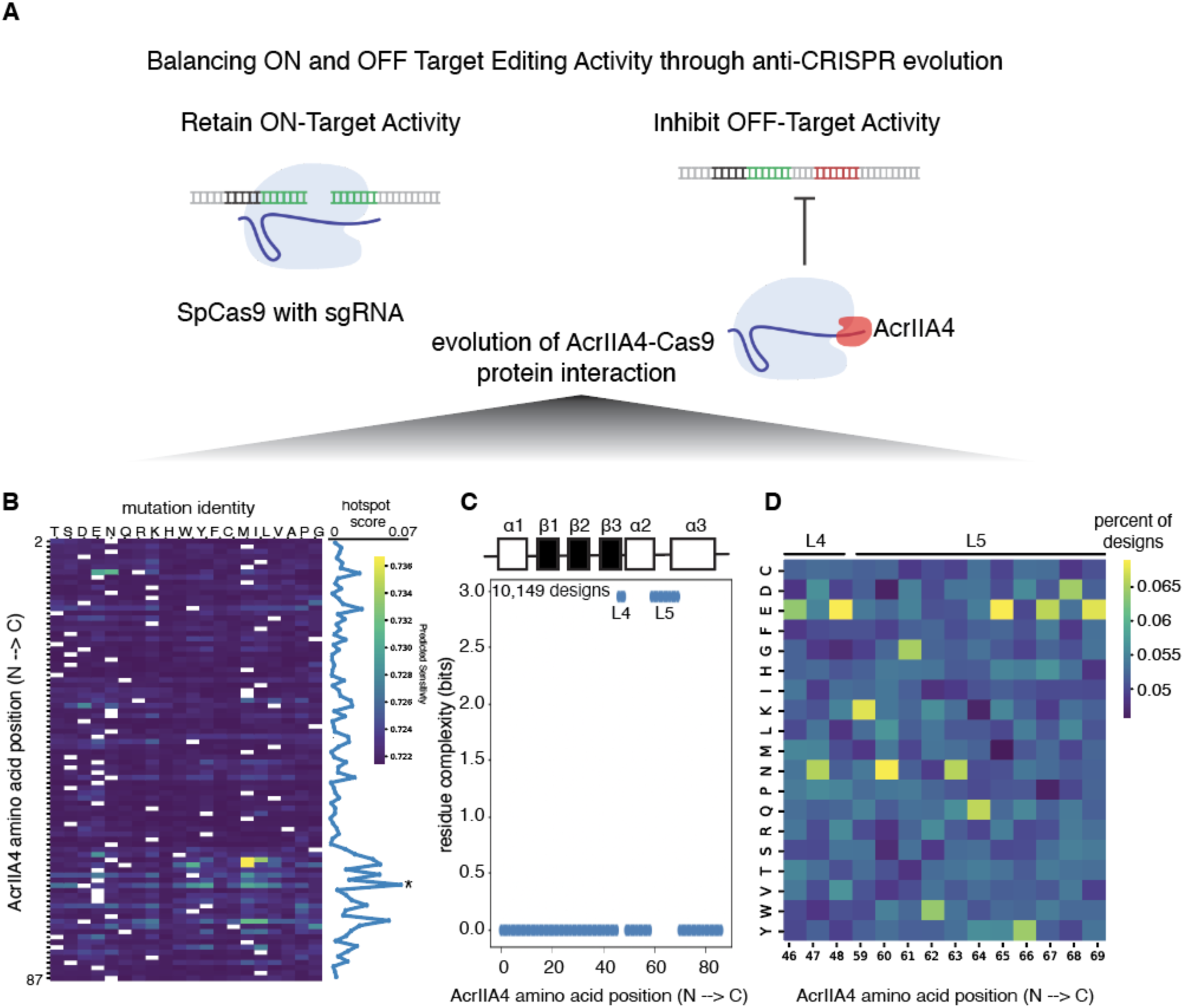
Protein language model-guided design of the AcrIIA4-Cas9 protein interaction. **A.** Schematic representing the goal of reducing off-target activity while retaining on-target activity through DNA-mimic anti-CRISPR protein engineering (AcrIIA4). **B.** Protein language model predictions of engineering hotspots within AcrIIA4. The matrix shows engineering hotspots (in yellow) that are predicted to be functionally significant. The sum of the hotspot score at each position is plotted to the right of the heatmap. **C.** Residue complexity at each amino acid position contained within the library of 10,149 designs. **D.** Frequency of amino acid identity (normalized to total designs in the library) at each position within the loop 4 (L4) and loop 5 (L5) regions.

Recently, protein language models have emerged as powerful tools to efficiently generate novel protein variants with altered activity. Protein language models (PLMs) learn underlying sequence constraints from multi-sequence alignments, which can be used as a representation of the underlying protein sequence-function relationships. Therefore, PLMs can be harnessed to compress the variant search space into an experimentally tractable set of variants that can be used to artificially engineer proteins with desired activity. We harnessed the ESM-1v PLM, which predicts the effect of protein sequence variation on function without requiring a model to be trained on each unique protein^22^.

The ESM-1v model is an ensemble model of 5 models that each compute the log-likelihood of observing a protein sequence variant given the sequence patterns within its training data. The uniformity of the ESM-1v predictions across the ensemble can be used as a proxy of evolutionary conservation that can capture long-range interactions that a standard multi-sequence alignment (MSA) cannot. We hypothesized that regions where most variants were predicted to have low probabilities would be functionally significant, having been selected due to their ability to inhibit CRISPR/Cas9-based immunity, thereby allowing the phage to propagate within a bacterial population. As a result, these regions could serve as engineering hotspots that, when mutated, would attenuate the ability of AcrIIA4 to fully inhibit SpCas9 activity. Using this approach, we constructed a hotspot score, which identified residues at the C-terminus of AcrIIA4 as a region that, if mutated, may attenuate AcrIIA4 function and tune SpCas9 activity (**Figure 1B, Methods**).

Since the structure of AcrIIA4 in complex with SpCas9 has been solved^18,23–25^, we executed a complementary rational design approach where we focused deep mutational scanning^26^ on loop regions (L4, L5) within the PLM-based hotspot that directly interact with SpCas9 to mimic its dsDNA substrate. In addition to deep mutational scanning, alanine scans were used to identify key residue sequence-function relationships. In total, the library was composed of 10,149 AcrIIA4 variants in which amino acids were uniformly represented within the engineering hotspot regions located in L4 and L5 **(Fig 1C, D**).

### Pooled protein screening to measure DNA repair outcomes across AcrIIA4 variants

Efficient engineering of AcrIIA4 within human cells requires a scalable and high-throughput assay to characterize gene editing precision over time. We took inspiration from conventional CRISPR screening methodologies, which group sgRNAs into lentiviral pools to query the resulting phenotypes of genetic perturbations at a genome-wide scale^4,5^. Here, instead of varying sgRNAs, we varied the AcrIIA4 amino acid sequence. We created a sequence-based reporter system (AcrobaTx barcode) that enables highly parallel profiling of on- and off-target editing across pooled AcrIIA4 variants. Briefly, the system contains both on- and off-target sites that are immediately downstream of the AcrIIA4 variant, thereby enabling measurement of DNA repair outcomes and Acr variant identity within a single sequencing read. The on- and associated off-target site were selected from the *AAVS1* locus (**Methods)**. Similar approaches have been developed to profile DNA repair outcomes across a pool of different sgRNAs^27–31^ and genetic perturbations^32^. The barcode was contained within a lentiviral vector together with a sgRNA targeting the on-target site and delivered to lung adenocarcinoma A549 cells expressing SpCas9 (**Figure 2A**). Therefore, sgRNA expression and RNP formation occurred in parallel with the expression of the AcrIIA4 variants within the pool.

**Figure 2.**
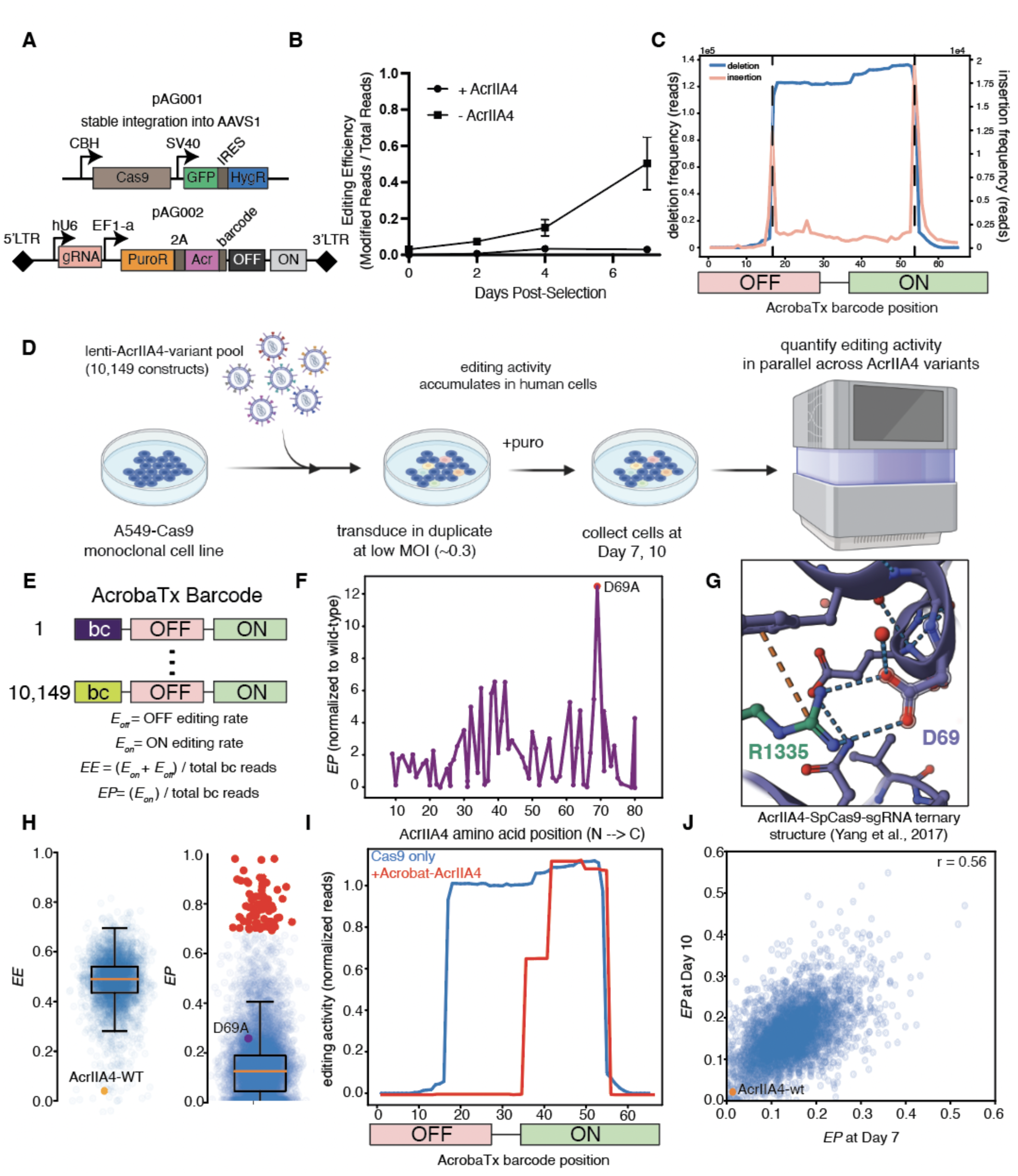
Pooled protein screening of AcrIIA4 variants in human cells. A. Two-component vector system where Cas9 is integrated into the AAVS1 safe harbor locus to sustain stable expression. The second component is a lentiviral system containing a sgRNA targeting on and off-target sites downstream of an AcrIIA4 variant that is uniquely barcoded. **B.** Editing efficiency measured over time within cells expressing wild-type AcrIIA4 (circular points) versus those without wild-type AcrIIA4 (rectangular). **C.** Indel mutation profiles spanning the on- and off-target regions. Deletions are shown in blue and insertions are in orange. **D.** Experimental design in which AcrIIA4 variants were delivered to A549 cells expressing Cas9, which were cultured for 10 days prior to collection and sequencing of the AcrobaTx barcode. **E.** Structure of the AcrobaTx barcode and definitions of gene editing activity metrics derived from measuring editing outcomes within the on- and off-target sites. **F.** Editing precision of alanine scanning variants normalized to the wild-type control. D69A variant is highlighted as the variant with the highest precision within the alanine scan mutant group. **G.** Crystal structure of AcrIIA4 bound to the SpCas9-gRNA complex. The interaction between D69 and R1335, required for PAM recognition, is highlighted. **H.** Distribution of editing efficiency (*EE*, left) and editing precision (*EP*, right) across all AcrIIA4 variants. Variants that resulted in *EP* in the top 90^th^ percentile are highlighted in red. **I.** Deletion profiles of DNA repair outcomes from cells without AcrIIA4 (blue) versus cells that express a variant that promotes precise editing (red). **J.** Editing precision measured across AcrIIA4 variants at Day 10 versus at Day 7 (Pearson Correlation Coefficient r = 0.56).

We reasoned that measuring DNA repair outcomes within the on- and off-target sites could effectively model the on and off-targeting activities of SpCas9 across each AcrIIA4 variant. To test this, we benchmarked the editing activity of Cas9 within cells that contained AcrIIA4 and those that did not. Measurement of DNA repair outcomes over time revealed accumulation of editing within ∼50% of cells in the absence of AcrIIA4 and inhibition of editing within cells that expressed AcrIIA4 (**Figure 2B**). DNA repair outcomes can be visualized within the barcode to distinguish on- and off-target editing events. Interestingly, frequent deletions were observed between the predicted cut sites in the on- and off-target regions, which contained three mismatches. The most frequent deletion was 38 nucleotides in length and exactly spanned the predicted cut sites, which is indicative of two separate editing events rather than microhomology-mediated end joining. In addition, insertions were observed at the predicted cut sites that are presumably representative of templated insertion events. Together, these two observations support the ability of the AcrobaTx barcode to model SpCas9 editing precision and suggest that SpCas9 promiscuously edited both the on- and off-target sites in the absence of AcrIIA4 **(Figure 2C)**. With the reporter system established, the first AcrIIA4 variant library, containing both alanine and deep mutational scans, was designed to interrogate whether systematic mutagenesis of the AcrIIA4-Cas9 interaction could result in tunable DNA repair outcomes.

In the initial pooled protein screen, A549 lung adenocarcinoma cells were transduced at low MOI (∼0.3) with a lentiviral pool of AcrIIA4 variants such that, on average, each cell contained only one variant. Low MOI transduction is important to avoid the confounding effect of multiple integrations, which could lead to variations in gene editing activity due to copy number and/or dominant negative effects rather than variant-specific effects. After transduction, cells were selected with puromycin and allowed to proliferate for 10 days. During proliferation, edits would accumulate within target regions that are differentiated by an upstream barcode that identifies the AcrIIA4 variant (**Figure 2D**). In other words, highly parallel measurements of DNA repair outcomes across AcrIIA4 variants enabled mapping of gene editing precision to AcrIIA4 variant identity. Across each variant, DNA repair outcomes were computed as either the proportion of all reads containing an edit at the on- and off-target sites (editing efficiency or *EE*), or the proportion of reads containing only on-target edits (editing precision or *EP*) (**Figure 2E**). As expected, the wild-type AcrIIA4 inhibited both on- and off-target editing, resulting in an *EE* of ∼0.05 (**Figure 2H, left**).

The alanine scans were first evaluated to determine the phenotypic consequence of removing R-group reactivity of a specific protein residue while keeping the protein’s backbone structure intact^33^. Focusing on the editing precision phenotype, we visualized the *EP* normalized to that observed within cells expressing wild-type AcrIIA4. Strikingly, a peak emerged at D69, which is a key residue that is known to directly interact with the PAM-interacting residue R1335 in SpCas9 and mediate binding affinity^23^ (**Figure 2F, G**). In addition, this residue was at a maximum within the hotspot region identified through the PLM-guided AcrIIA4 variant library design process (**Figure 1B, asterisk**). Together, these results suggest that ablating the D69 electrostatic interaction with R1335, which reduces AcrIIA4 binding affinity to SpCas9, increases gene editing precision in A549 human cells.

While the alanine scanning results supported known interactions between AcrIIA4 and SpCas9, we sought to identify novel variants that enhance gene editing precision through deep mutational scanning. First, we calculated the *EE* and *EP* for every variant. First, the median *EE* was ∼0.5, which is consistent with the overall editing efficiency calculated in cells that did not express AcrIIA4 (**Figure 2B, H**). Second, we observed frequent editing at the off-target site across variants. However, a subset of variants (above the 90^th^ percentile of *EP*) showed nearly complete depletion of off-target editing activity (**Figure 2H, right**). Third, analysis of the indel spectrum within a top-ranked variant (enAcr-1) revealed editing activity that was restricted to the on-target site, suggesting that the AcrIIA4 variant was sufficient to inhibit off-target editing activity within this pooled readout (**Figure 2I**).

Finally, we examined the durability of the *EP* phenotype over time by measuring DNA repair outcomes at both Day 7 and Day 10. We observed that the AcrIIA4-WT stably inhibited editing activity and that there was a reasonable correlation in *EP* across both time points for all variants (PCC = 0.56) (**Figure 2J**). Notably, this correlation was sensitive to the sampling depth (number of sequencing reads) across all variants. To address this sensitivity, we introduced a clonal tag in subsequent experiments to measure the numbeer of different cellular populations that express each variant (**Methods**). The clonal tag provides additional information to directly address potential confounding effects whereby *EP* is driven by clonal bottleneck effects. From a technical perspective, these results support the accuracy of our technology to deconvolute DNA repair outcomes across AcrIIA4 variants that are expressed in a pooled fashion across millions of cells and motivate further reproducibility and validation studies.

### Validation screen to identify lead AcrIIA4 variants that increase gene editing precision in human cells

Variants were rank ordered by *EP* and the top two thousand were resynthesized into a new variant validation pool for testing within an expanded cell line panel that included A549 that expressed Cas9 (A549-C9) and 293T human embryonic kidney cells expressing either Cas9 (293T-C9) or high-fidelity Cas9-HF1^15^ (293T-C9-HF1). The same experimental design was used to measure *EP* after 7 days of editing as in round 1 (**Figure 3A**). Within A549 cell lines, the median of the *EP* distribution increased from 0.25 in round 1 to 0.55 in round 2 across all variants. Interestingly, the median of the *EP* distribution was higher within the 293T-C9 context suggesting that the cell line context, potentially due to differences in Cas9 activity, sgRNA expression levels or DNA repair pathway activity, can influence the *EP* distribution (**Figure 3B**). Since the Cas9 and sgRNA constructs are conserved across cell lines, the differences may be biological and reflective of the underlying DNA repair pathway activity or p53 status^34^. Lastly, the *EP* distribution reflecting editing within 293T-C9-HF1 cells had a significantly higher median of 0.94 relative to all other sample groups. Consistent with expectations, this suggests that Cas9-HF1 had increased targeting specificity relative to SpCas9.

**Figure 3.**
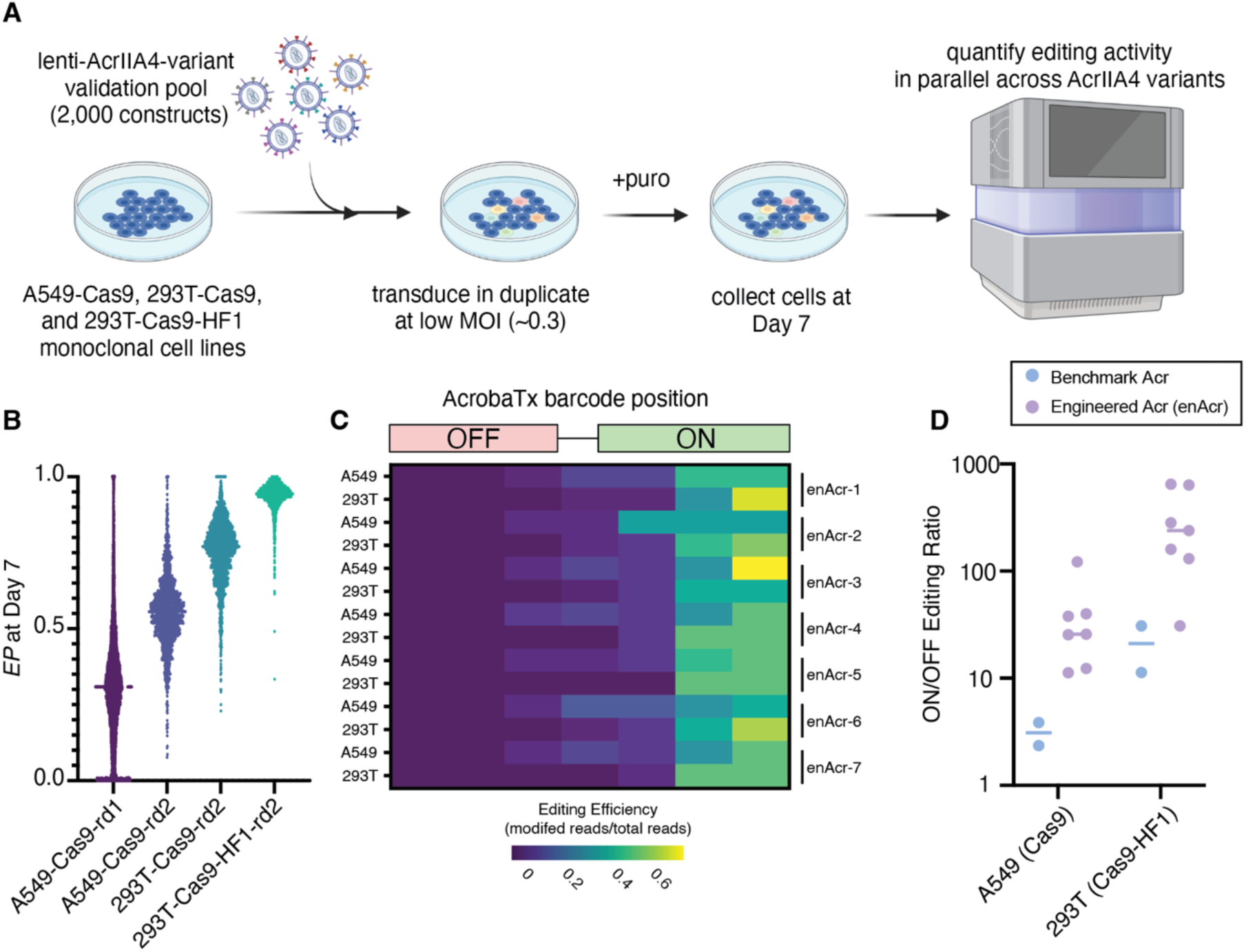
Validation screen to assess reproducibility of AcrIIA4-driven editing precision. **A.** Experimental design in which the top 2000 AcrIIA4 variants, rank-ordered by *EP*, were delivered to A549 and 293T cells for another round of screening. **B.** Distribution of *EP* across all sample groups in the second round of screening versus the first round, at left. **C.** Frequency of editing within the AcrobaTx barcode across seven lead candidate enAcr groups. Yellow rectangles indicate areas in which editing occurs frequently while violet depicts areas with low editing activity. **D.** Ratio of on- to off-target editing in cells expressing seven representative lead candidate enAcrs (purple) versus ratios observed from DNA repair outcomes within cells expressing previously described benchmark AcrIIA4 variants.^19^

Seven lead AcrIIA4 variant candidates were identified based on their validation screen *EP* values (**Methods**). Within these seven variants, there was an enrichment of hydrophobic residues in L5 which, in the wild-type form, contains hydrophilic residues that form electrostatic interactions with the CTD of SpCas9^25^. Their editing activity was visualized within the AcrobaTx barcode. Across all variants, minimal off-target editing was observed (<1%) whereas clear on-target editing (∼60%) was achieved. There were no detectable off-target edits within the 293T-C9-HF1 cell line context (**Figure 3C)**. To further assess the benefit of the engineered AcrIIA4 variants (enAcr-1-7), the ratio of on-target editing to off-target editing was calculated and compared to published benchmarks that were reported to increase gene editing precision^19^.

Strikingly, there was a significant increase in the on-target to off-target editing ratio within the enAcr group relative to the benchmarks and also an increase in the 293T-C9-HF1 setting (**Figure 3D**). Together, these results suggest that potentially concurrent delivery or pre-delivery of enAcrs could significantly reduce off-target editing activity across cell line contexts and in synergy with Cas9-HF1.

### Engineered AcrIIA4 variants diminish off-target editing activity while retaining on-target editing activity

Cas9 RNP-based delivery has been implemented for editing in primary human cell contexts^35,36^. While the precision of editing with RNPs is generally higher than viral-based delivery, off-target events are still observed and can lead to aneuploidy^37,38^. We hypothesized that the enAcr molecules could serve as a prophylactic to reduce off-target activity in cells that are edited by SpCas9 in its RNP format. An advantage of this approach is that it does not require re-dosing of a wild-type AcrIIA4 (AcrIIA4-AT), which has been shown to inhibit on-target editing to a lesser extent (∼2-fold) than off-target editing (∼4-fold)^18^. To test this, we generated 293T cell lines that stably expressed enAcr-1, AcrIIA4-WT, and AcrVA1. AcrVA1 does not bind to SpCas9 and therefore served as negative control.

We delivered RNP complexes through electroporation to cells that were individually transduced with the anti-CRISPR panel of proteins. After 96 hours, we measured on- and off-target editing in cells that contain a gRNA that targeted beta-globin (*HBB*) or *AAVS1.* The latter was the target sequence used in the AcrobaTx barcode (**Figure 4A**). Importantly, beta-globin is an independent locus that is both therapeutically relevant and contains numerous off-target sites.

**Figure 4.**
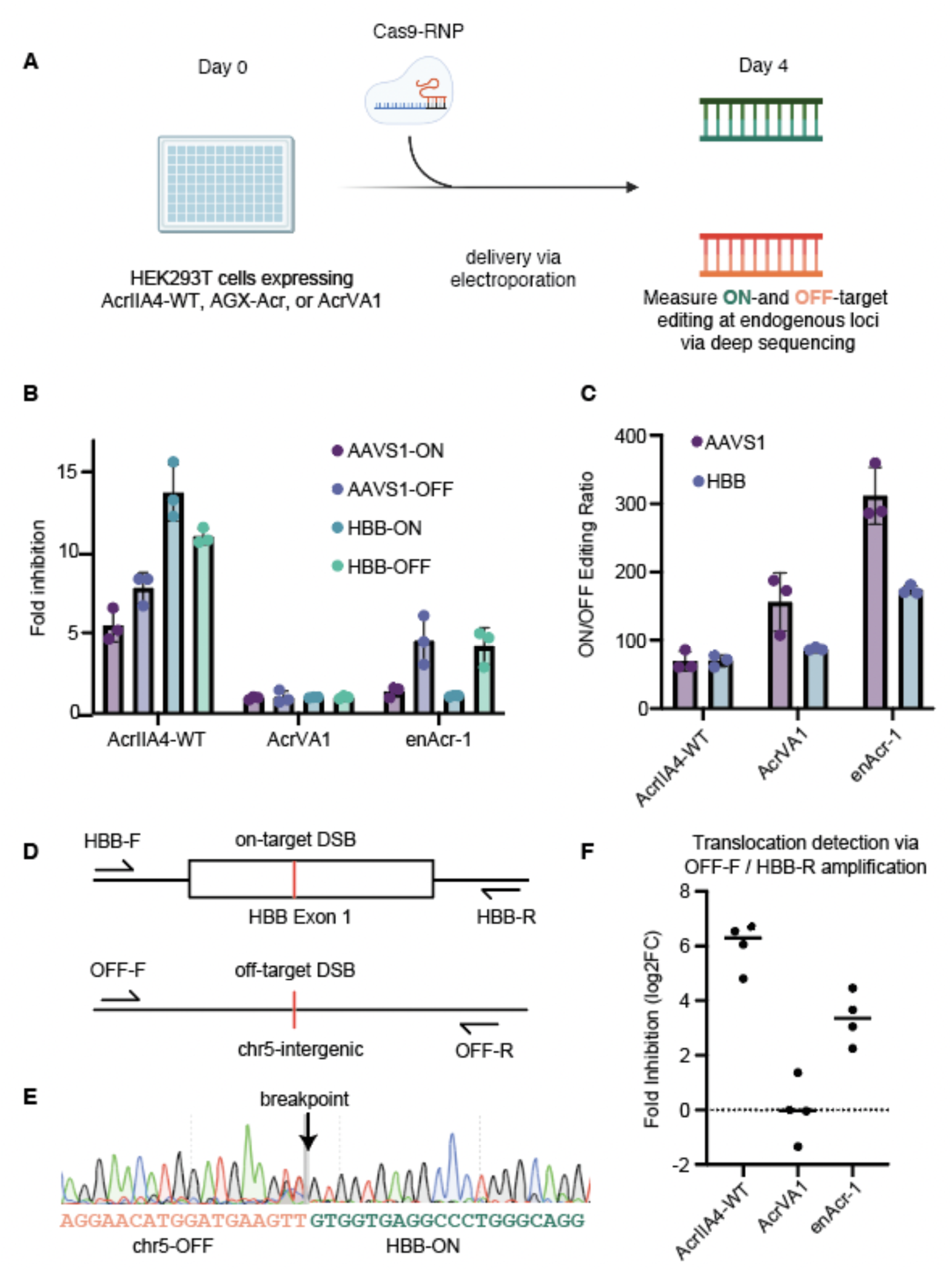
Differential inhibition of on- and off-target editing activity in cells expressing engineered AcrIIA4 variants. A. Experimental design where cells stably expressing anti-CRISPR proteins were electroporated with Cas9-RNP and incubated for four days prior to measurement of on- and off-target editing activity through deep sequencing. **B.** Engineered AcrIIA4 variant (enAcr-1) diminishes off-target editing activity while retaining on-target activity at the *AAVS1* and *HBB* loci. The wild-type AcrIIA4 diminishes both on- and off-target editing activity. **C.** Ratio of on- and off-target editing at the *AAVS1* locus in cells expressing AcrIIA4-WT, AcrVA1, or an engineered AcrIIA4 variant (enAcr-1). **D.** Design of PCR-based assay to detect chromosomal translocations between *HBB* and an intergenic off-target region located on chromosome 5. **E.** Sanger sequencing identifies a breakpoint at the predicted cut site within the off-target region (orange) that is then ligated to HBB (green). **F**. qPCR-based quantification of translocation events from cells expressing AcrIIA4-WT, AcrVA1, or enAcr-1. Quantifications are normalized to the average amplification level measured in cells expressing the AcrVA1 control.

Consistent with the pooled screening experiments, the ratio of on- to off-target editing at the *AAVS1* and *HBB* loci increased by approximately 2-fold in cells expressing enAcr-1 relative to cells expressing the inert AcrVA1 control protein **(Figure 4C**). As expected, the AcrIIA4-WT inhibited both on- and off-target editing by 5.4-fold and 7.8-fold, respectively, when compared to the AcrVA1 control at the *AAVS1* locus (**Figure 4B**). In agreement, there was a consistent overall reduction of editing activity at both the on- and off-target sites of *HBB* in cells expressing AcrIIA4-WT (**Figure 4B**). Notably, the enAcr-1 simultaneously inhibited off-target editing by approximately 4.5-fold while retaining on-target editing at both loci (**Figure 4B**).

We sought to identify whether a translocation occurred between *HBB* and an off-target region located on chromosome 5 (chr5-OFF) using a PCR-based approach since they have been previously observed^52^. We identified a novel translocation event that resulted in the ligation of cleaved *HBB* downstream of the cut site located in the off-target region (**Figure 4D,E**). We hypothesized that there would be a reduction in translocation events in cells expressing enAcr-1 since it is able to inhibit off-target editing that is required for translocation to occur. We quantified the abundance of translocation events within a representative sample of cells and found that, relative to cells expressing AcrVA1, cells expressing enAcr-1 reduced the abundance of translocation events by about ∼9.8 fold (**Figure 4F, Methods**). Together, these results suggest that the engineered AcrIIA4 variants can increase the specificity of editing, by selectively diminishing off-target editing activity, across endogenous loci within the human genome.

## DISCUSSION

Ensuring efficient targeting while minimizing undesired genome modifications will become increasingly important as gene editing therapies become deployed more widely in clinical settings^6,7,39,40^. Here we demonstrate an integrated computational and experimental platform that can generate protein candidates and simultaneously measure on- and off-target editing activity for ten thousand or more variants at once. This approach adds to the expanding repertoire of methods available to perform deep mutational scans in mammalian cells and builds upon previous platforms that have been developed to screen anti-CRISPR proteins within cell-free settings^53,54^. Using this platform, we engineered novel AcrIIA4 variants that maintain desired on-target editing while significantly reducing off-target editing events. Extensive work has been done to develop small-molecule inhibitors to tune SpCas9 activity^41,42^. This work adds to this pre-existing toolkit with components that can seamlessly be incorporated into existing recombinant gene therapy vectors that contain SpCas9. In addition, the AcrIIA4 variants can be co-expressed with AcrIIA4-WT to enable tissue-specific genome editing^43,44^ and to design genetic circuits to control cellular behaviors^45^.

Anti-CRISPR molecules serve as attractive targets for engineering substrates for tuning CRISPR activity due to their compact size and evolved specificity of activity^49^. In addition, they form a separable functional domain that can be engineered orthogonally from CRISPR systems. Furthermore, given the growing diversity of identified anti-CRISPRs, we anticipate that our approach will be easily extensible to a wide range of Cas effectors. Finally, the AcrobaTx platform is agnostic to a particular application because it uses a protein language model that performs zero-shot predictions and a high-throughput screening approach that can be generalized to a wide array of protein functions^22^. We anticipate that this approach will help to generate custom proteins with defined activities, which will accelerate the development of technologies in the gene editing field and beyond.

## LIMITATIONS OF THE STUDY

Our study does not determine the details of how the novel AcrIIA4 variants modulate editing precision. We speculate a few mechanisms centered around modulation of AcrIIA4-SpCas9 binding could be at play. First, a reduction in AcrIIA4 binding affinity to SpCas9 may be sufficient to switch between almost total inhibition of SpCas9 to releasing a small number of cleavage-competent molecules, which may kinetically bind preferentially to and cleave the on-target locus^19^. Second, mutagenesis of the contact of the AcrIIA4 and the CTD of Cas9 may allow for initial PAM recognition followed by R-loop formation. The remaining interactions between the AcrIIA4 variant and the SpCas9 TOPO and RuvC domains may trap the complex in an inactive state and prevent nuclease activity. However, full complementarity between the sgRNA and target DNA may provide sufficient binding energy and torque to drive off variant AcrIIA4 while the torque generated by partial R-loop formation resulting from sgRNA-DNA mismatches may not^46–48^ (**Figure 5**). Further investigation will help to elucidate the mechanism of precision enhancement and contribute to a more detailed understanding of Cas9 and AcrIIA4 function.

**Figure 5.**
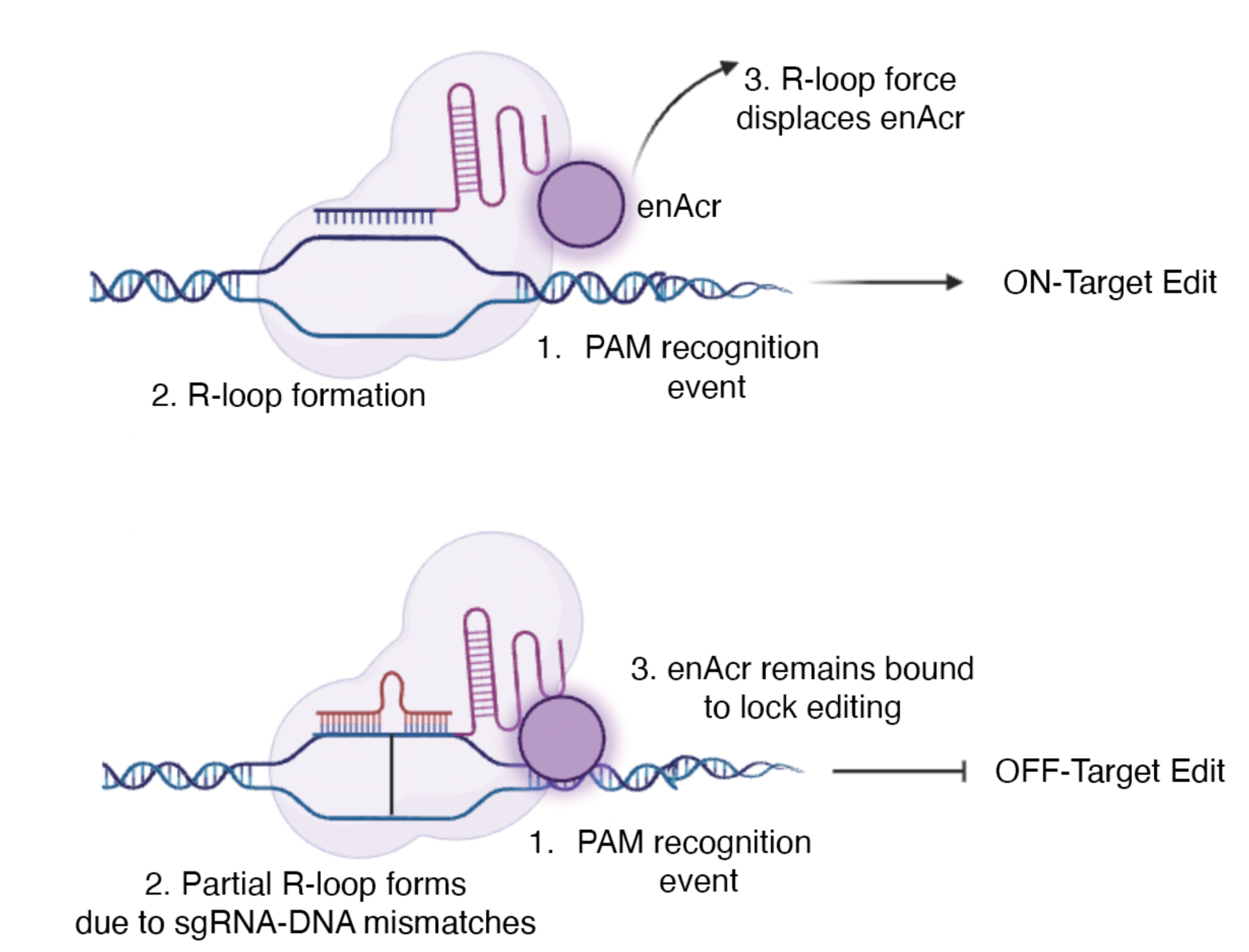
Proposed model for differential inhibition of on- and off-target editing activity through engineering of the interaction between AcrIIA4 and Cas9.

## SUPPLEMENTARY MATERIALS

### Methods

#### Computational identification of protein engineering hotspots

All possible single amino acid mutations were generated across the reading frame of AcrIIA4 (20x87 matrix) and used as input into the ESM-1v model. Each of the 1740 single substitution variants had an associated ensemble of log-likelihoods generated by the model. The uniformity of the ESM-1v ensemble prediction distribution was quantified using the Shannon entropy metric. The inverse of the entropy metric was computed to generate a hotspot score, in which higher values are predicted to have functional significance, and were plotted across all variants to identify engineering hotspots (**Figure 1B**).

#### AcrIIA4 primary screen library design

After identification of hotspots through an unbiased ESM-1v-guided approach, empirical structural information was used to further refine the hotspot based on AcrIIA4 contact with SpCas9. A custom python script was used to randomly generate amino acid sequences within the engineering hotspot and to reverse translate the variant amino acid sequence into a human codon optimized DNA sequence space. The DNA sequences were further modified to include cloning sites and unique barcode sequences for identification prior to assembly within a lentiviral backbone.

#### Molecular cloning of the AcrIIA4 benchmark lentiviral backbone

A lentiviral backbone was cloned in which typeII-S cloning sites were inserted downstream of a puromycin resistance cassette that contained a 2a peptide to enable in-frame ligation of AcrIIA4 variants in a pooled fashion. Additional artificial SpCas9 target sites were cloned downstream of the type II-S cloning sites through the NEB HiFi assembly approach using the manufacturer’s instructions to produce the final pAG002 backbone. To benchmark the ability of the SpCas9 synthetic target sites to report AcrIIA4 activity, wild-type AcrIIA4 was cloned downstream of the 2a peptide along with an empty negative control that contained the parental typeII-S cloning site sequence.

For the second validation screen, a new lentiviral backbone was constructed that introduced a clonal tag into the AcrobaTx barcode. Briefly, an oligonucleotide (IDT) containing a 10 base pair random sequence was ordered and inserted through the NEB HiFi assembly approach using the manufacturer’s instructions. The representation of the 10 bp random sequence was measured using a MiSeq (Illumina).

#### Lentiviral preparation and delivery to benchmark AcrIIA4 activity

Lentivirus was prepared by co-transfecting pAG002 with VSV-G envelope and Delta-Vpr packaging plasmids into HEK-293T cells using Lipofectamine 3000 reagent (ThermoFisher). Supernatant was harvested 48 hours and 72 hours after transfection. A549 cells expressing the pAG001 Cas9 vector (Genecopeia) were transduced at high MOI with 8 μg/mL polybrene using a spin-infection at 1200*g for 45 minutes. After 48 hours, cells were selected with 1 μg/mL puromycin to establish stably expressing cell lines for gene editing analysis.

#### Large-scale lentiviral preparation and pooled screen tissue culture

Lentivirus was prepared as described above with slight modifications. First, lentivirus was concentrated by 10X using PEG-it virus precipitation solution (System Biosciences) according to the manufacturer’s instructions. Concentrated virus was then used for a functional titer in which a concentration consistent with an MOI of ∼0.3 was identified. Subsequently, cells stably expressing SpCas9 (pAG001) were transduced at a scale to achieve ∼1000X cells / Acr variant after puromycin selection of the pAG002 pooled library component.

#### Next-generation sequencing and analysis workflow

Primers were designed to amplify the AcrobaTx cassette containing the Acr variant-specific barcode sequence and Cas9 target sites. The resulting libraries were sequenced on a NovaSeq 6000 (Illumina).This cassette was used to measure editing efficiency at both the on- and off-target sites to derive metrics for overall editing efficiency (*EE*) and editing precision (*EP*). Reads were processed using a custom pipeline. Briefly, reads were trimmed and merged using publicly available software (cutadapt and FLASh). Next, the target region was assigned to an Acr variant using the variant’s unique barcode. When available, the clonal tag was error corrected to group tags that were within a hamming distance of 2 together, thereby allowing an assignment of sequence reads into clonal groups. The target region was then aligned to the reference sequence to call indel mutations using the Needleman-Wunsch algorithm^50^. Finally, the alignments were parsed to identify mutations within the predicted target window (three base pairs upstream of the PAM site).

#### Selection of lead engineered AcrIIA4 variants

A data frame was constructed using the validation screen data set that included the editing precision, editing efficiency, and clone number (based on the clonal tag) for each AcrIIA4 variant. First, all AcrIIA4 variants were removed that were represented by fewer than 20 independent clonal populations. Next, AcrIIA4 variants that were in the bottom 50^th^ percentile of the editing efficiency distribution were removed. The variants were then rank-ordered by editing precision and the top 7 were selected for further validation studies.

#### Ribonucleoprotein delivery of SpCas9

The electroporation procedure was conducted using the InvitrogenTM NeonTM Transfection System with 100 μL tips. Alt-R® CRISPR-Cas9 crRNA and Alt-R® CRISPR-Cas9 crRNA XT (IDT) were reconstituted in IDTE Buffer to achieve a final concentration of 200 μM each.

Subsequently, these oligos were mixed in equimolar proportions in a sterile microcentrifuge tube, resulting in a final duplex concentration of 100 μM. The duplex complex underwent denaturation at 95°C for 5 minutes and was then allowed to cool at room temperature for 5 minutes.

For the formation of the ribonucleoprotein (RNP) complex, 120 pmol of each Alt-R guide RNA was combined with 104 pmol Alt-R™ S.p. Cas9 Nuclease V3 and incubated at room temperature for 15 minutes. Cell lines stably expressing anti-CRISPR proteins were trypsinized, washed twice with PBS, and resuspended in 100 μL of resuspension buffer R plus the RNP complex. Electroporation was carried out using the following program: Pulse voltage: 1,100 V, Pulse width: 20 ms, and Pulse number: 2. Post-transfection, cells were plated in prewarmed medium and collected 4 days after transfection for subsequent analysis.

#### Quantification of editing at endogenous genomic loci

Primers were designed to amplify the targeted endogenous genomic loci. The resulting libraries were sequenced on a MiSeq (Illumina). Raw reads in FASTQ format were analyzed for editing activity at each on- and off-target site by CRISPREsso2^51^ using the batch settings and default parameters for Cas9: Center of the quantification window -3, Quantification window size of 2 base pairs with substitutions excluded. The median number of reads across all samples was approximately 100,000 reads per sample. Fold inhibition in editing was determined using quantification of editing within the AcrVA1 as the control.

#### Quantification of translocation events

qPCR was conducted with PowerUp™ SYBR™ Green Master Mix for qPCR (Applied Biosystem) in a 10 μl reaction volume on a QuantStudio™ 7 Flex Real-Time PCR System (Applied Biosystem) using 50 ng of DNA per reaction. The standard cycling mode as suggested by the manufacturer was used: 50oC for 2 minutes, 95oC for 2 minutes and 40 cycles of 95oC for 15 seconds (denature) and 60oC for 1 minute (annealing). Image acquisition was performed on each cycle after the annealing step. A melt curve was performed after the PCR for quality control. The relative quantification was achieved using ΔΔCq analyses with *GAPDH* as the reference gene and levels within the AcrVA1 cells as the control.

## ACKNOWLEDGEMENTS

The authors thank Dr. Doug Fowler, Dr. Brian Hie, and Dr. Monte Winslow for comments on this study and manuscript. In addition, the authors thank Cynthia Kuan for operational support. The authors also thank Devyani Jogran, Shalaka Wahane, Marc Martin-Casas, and Amanda Cashin for laboratory support with next-generation sequencing.

## AUTHOR CONTRIBUTIONS

Methodology, J.M. and N.W.H.; Formal Analysis, J.M. and N.W.H.; Investigation, J.M., K.V., E.K., and N.W.H.; Resources, L.C.; Writing, J.M., M.N., L.C., and N.W.H., Funding Acquisition, N.W.H.

